# Electroacupuncture regulates microglia activation through the STING/NF-*κ*B pathway to reduce pain in bone cancer pain model

**DOI:** 10.1101/2024.05.08.593142

**Authors:** Ying Liang, Wenhao Liu, Zhiyi Shen, Xu Yan, Yihan He, Nenggui Xu

## Abstract

Cancer pain is a global public health problem. The mechanism of cancer pain is complex, and opioid analgesics, which are widely used clinically, have obvious addiction and side effects, which seriously affect patients’life functions and may aggravate their anxiety, depression and other negative emotions. Acupuncture has a history of thousands of years in China, and acupuncture analgesia has been confirmed by many studies. This study investigated whether electroacupuncture can alleviate abnormal pain in bone cancer pain (BCP) mouse models and its possible central mechanism. A bone cancer pain model was established by injecting Lewis lung cancer cells into the left femoral cavity of adult male mice. Mechanical paw withdrawal threshold was tested baseline before surgery and 1, 4, 7, 10, 14 and 21 days after surgery. On day 21, behaviours related to depression emotions were tested. After the behaviours, the femurs were removed to observe pathological changes, the neck was broken and brain tissue was collected from the basal lateral amygdala (BLA) area for subsequent Western Blot and ELISA experiments were performed to verify the expression of (stimulator of interferon genes, STING) STING/NF-*κ*B pathway proteins and the expression of inflammatory factors. Immunofluorescence of Ionized calcium-binding adapter molecule-1 (Iba-1) and STING in the basal lateral amygdala (BLA) brain region was also performed. The results show that electroacupuncture can increase the pain threshold of the bone cancer pain model and alleviate the depressive-like emotional phenotype. Electroacupuncture inhibited the expression of STING/NF-*κ*B pathway proteins, activation of microglia and release of inflammatory factors in the basal lateral amygdala (BLA) area. Therefore, this study shows that electroacupuncture may relieve bone cancer pain by regulating microglial activation and inflammatory factor release through the STING/NF-*κ*B pathway.

## Introduction

Bone cancer pain is currently a challenging research topic in the field of neurobiology and lacks effective treatment options [1]. Pain is associated with microglia activation and increased inflammatory factors in the brain [2]. Depression is often comorbid with malignancy and can further aggravate the original disease [3]. Pain messages are transmitted through the peripheral spinal cord to different areas of the brain, and the BLA is the most important brain region [4].

Microglia play an active role in brain regions important for emotional and memory-related aspects of chronic pain [5]. Chronic stimulation is associated with microglia activation and neuroinflammation [6]. Following activation, microglia can adopt pro-inflammatory M1 and anti-inflammatory M2 phenotypes. The M1 phenotype of microglia can induce an increase in protein expression, such as inducible nitric oxide synthase (iNOS) and cluster of differentiation Arginase-1 (Arg-1), and promote the release of pro-inflammatory cytokines [7].

Electroacupuncture therapy has been widely used in clinical practice, and currently acupuncture for cancer pain is gradually becoming the NCCN Cancer Treatment Guidelines, an oncology clinical practice application guideline issued by the National Comprehensive Cancer Network (NCCN) in the United States of America, and has achieved good analgesic efficacy [8]. Acupuncture analgesia is a combined effect in which signals generated by acupuncture points are transmitted to relevant areas of the spinal cord and brain, increasing or decreasing a variety of neurotransmitters, inflammatory factors to reduce pain [9]. Foot Sanli is a commonly used acupoint for analgesia in clinical practice [48]. Animal studies have shown that the combination of Foot Sanli and Sanyinjiao is a superior acupoint combination for the treatment of painful depressive comorbidities [10].

In this study, we induced bone cancer pain in WT mice after inoculation of Lewis lung cancer cells (LLC) into the femoral intramedullary canal. We found that BCP mice showed a significant decrease in pain threshold and sugar-water consumption compared to sham mice. Electroacupuncture reduced mechanical pain sensitivity and depression. Mechanistically, electroacupuncture attenuates bone cancer pain and depressed mood by inhibiting the STING/NF-*κ*B signalling pathway, microglia activation and inflammatory factor release.

## Materials and methods

### Animals

Male SPF C57BL/6 mice (17-19 g, n = 30) were obtained from Guanggdong Zhiyuan Biomedical Technology Co. (Guangzhou, Guangdong). Animals were placed in cages and allowed to rest for one week under controlled environmental conditions (23±2ºC, 12 h light/12 h dark) to acclimate to the new environment. They were divided into three groups: SHAM SURGERY, MODEL, MODEL+EA groups.

### BCP model

Lewis lung cancer (LLC) cells were digested with 0.05% trypsin, and a suspension of 5×10^7^/mL cells was made in PBS. Mice were injected with 0.3 ml of aflatoxin solution, and when the mice were anaesthetised, they were fixed on an operating table, hair near the femoral plateau of the mice was shaved, a towel was spread, and a 0.5-1 cm superficial incision was made near the femoral plateau to expose the patellar ligament. A 25-gauge needle was inserted into the femoral cavity at the left intercondylar notch of the femur, and then 2×10^5^ cells were implanted with a 10 *µ*L microinjector. To prevent leakage of tumour cells outside the bone cavity, the injection port was sealed with silicone glue at the site of the external injection, the wound was closed, and penicillin sodium was applied to the incision. For the sham-operated control group, the steps were the same as above, and PBS solution without tumour cells was injected in the femoral cavity [11].

### Electroacupuncture

The acupuncture needles were sterile disposable acupuncture needles, 0.18 mm × 10 mm. Acupuncture points are selected according to the method in ≪Chinese Veterinary Acupuncture≫ [12]. With reference to the positions of relevant acupuncture points in the human body, they are positioned using the method of animal comparative studies; Zusanli (ST36); Sanyinjiao (PC6); two points, 3mm in Zusanli and 1.5mm in Sanyinjiao. After the acupuncture is completed, connect the power supply of the electroacupuncture device to the needle handle, connect the Korean acupuncture nerve stimulator (HANS-200A), and stimulate The parameters are: continuous wave, 2Hz, 1mA, 20min. Electroacupuncture treatment was started 10 days after modelling, once a day. Animals in the sham surgery group were injected with PBS only. All animals were used for emotional behaviour tests and brain tissue was collected for further experiments. Four animals from each group were used for histological analysis including haematoxylin and eosin staining and immunofluorescence staining. Four animals from each group were used for ELISA and Western blot analyses. The experimental procedures were performed in strict accordance with the relevant regulations of the “Guiding Opinions on the Kind Treatment of Experimental Animals” issued by the Ministry of Science and Technology of the People’s Republic of China. The animal experiment protocol was reviewed and approved by the Animal Ethics Committee of Guangzhou University of Traditional Chinese Medicine, Ethics No: GZY20221201003.

### Paw withdrawal mechaical threshold

The mechanical withdrawal threshold was determined by vonFrey (Ugo Basile, Italy) as previously described [13]. Mice were habituated to the environment for 30 min. After the mice were quiet, the fibres was aligned with the centre of the left plantar foot, the force was applied evenly to keep the fibres in a the fibres in a “C” shape for 5 s, and the mice were considered positive if they withdrew their feet. The test was performed sequentially with 0.16 g, 0.4 g, 0.6 g, 1.0 g, 1.4 g and 2.0 g of fibres, and each gram was repeated 5 times with an interval of of 1 min. If the same gram was positive 3 times, the gram was the mechanical threshold for the mice, and if it was negative 3 times, the next gram of fibres was used for the test. We tested mechanical thresholds in mice using von Frey on days -1, 0, 1, 3, 5, 7, 10 and 14.

#### Source preference

Prior to the experiment, a 24 hours sugar water acclimation trial was conducted 0.1% sugar water was placed in each cage with an equal amount of pure water, and the position of the water bottles was changed every 12 hours to prevent positional preference, and the mice were fasted from water and food for 12 hours prior to the formal experiment, and the sugar water consumption of the mice was calculated over the 24 hours period.

#### Western blot

BLA brain tissue was homogenised and centrifuged (15,000 rpm, 45 min) in RIPA lysate (Beyotime, Shanghai, China) and protease inhibitor cocktail (APExBIO, USA) and phosphatase inhibitor (APExBIO, USA) and the supernatant was collected. Sample concentration BCA protein assay kit (Beyotime, Shanghai, China) was used. Proteins were denatured and heated in a water bath at 95°C for 10 min. Protein samples were separated on 10% glycine SDS-PAGE gels and electrophoresed at 80 V for 30 min, then adjusted to 110 V for 60 min. Proteins were transferred to a PVDF membrane using the Trans-Blot Turbo system (Bio-Rad Laboratories, USA). The membrane was washed with TBST solution and then blocked with Quick Blocking Buffer (NCM biotech, Suzhou,China) for 30 min at room temperature. After washing for 5 minutes 3 times, the membrane was immersed in primary antibodies, including anti-STING (1:1000, rabbit, CST, 504945S), anti-P-TBK1 (1:1000, rabbit, CST, 5483T), anti-TBK1 (1:1000, rabbit, CST, 38066S) and other primary antibodies, anti-NF-*κ*B (1:1000, rabbit, Abcam, AB32536), anti-P-NF-*κ*B (1:1000, rabbit, CST, 3033T) and anti-GAPDH (1:4000, rabbit, Abways, AB0037) and stored overnight in a refrigerator at 4°C. Finally, after washing, the membrane was incubated with sheep anti-rabbit HRP-conjugated secondary antibody (1:30,000, Abcam, ab6721, rabbit) for 2 hours at room temperature. (The Western blots were then visualised using People’s Chemiluminescence System (ECL) in luminescence solution (Meilun Bio, China). Chemiluminescence System (Peiqing Technology Co., Ltd., China) for observation.

### Immunofluorescence

C57BL/6 mice were selected for modelling and grouping, and after acupuncture intervention, mice were anaesthetised by intraperitoneal injection of 0.4 ml Alfontaine solution, mice were fixed on the operating table, and the heart was successively perfused with 300 ml of 0.9% NaCL solution and 200 ml of 4% PFA solution, and then the head was cut off and the brain was removed, and the removed brain tissues were immersed in 4% paraformaldehyde and fixed for 24 h, and then 15%. The brain tissues were immersed in 4% paraformaldehyde and fixed for 24 h. The brain tissues were then dehydrated by gradient dehydration with 15% and 30% sucrose solution so that the brain tissues could be completely glycolised. The cut brain slices were rinsed three times with PBS for 10 min each time to clean the residual OCT; the sealing solution was added and the slices were sealed in a 37°C water bath for 1-2 h; after sealing, the brain slices were added with the primary antibody Iba-1 (abcam, ab289874, USA) dilution solution and left at 4°C overnight, and then rinsed three times with the PBS solution for 10 min each time; the fluorescent secondary antibody (Rabbit anti-Goat IgG HL(Alex488, AB150141) dilution solution was added, and the slices were incubated at 37°C for 1-2 hours, and then rinsed with the PBS solution for 10 min each time for three times; the brain slices were then frozen with a freezing section machine. The brain slices were then rinsed with the PBS solution for 10 min each time for three times, mounted on slides, stained with DAPI (beyotime, China) nuclear stain, sealed with 50% glycerol, stored at 4°C away from light, and observed and photographed with a laser confocal microscope.

### Enzyme linked immunosorbent assay

BLA brain tissue was homogenised in PBS, centrifuged and the supernatant extracted. The protein concentration of the sample was determined using the BCA Protein Assay Kit (Beyotime, Shanghai, China). The levels of IL-1*β* in BLA and serum were determined by ELISA kit (MeiMian, China), and the operation was carried out in strict according to the instructions.

### Statistical Analysis

The data in the experimental were expressed by mean*±*SEM, and if the data satisfed normal distribution and variance chi-square, independent sample t-test was used for comparison between two groups; for comparison of three groups and above, one-way ANOVA test was used, and if variance chi-square was satisfed, LSD test or Bonferroni correction was used for two-way comparison. If variance chi-square was not satisfed, Tamhane test was used the nonparametric Kruskal–Wallis test was used for multiple group comparisons. All experimental data were statistically analysed by GraphPad Prism 9.0, and P*<*0.05 was considered a statistically signifcant diference between groups. Plotting was performed by GraphPad Prism 9.0.

## Results

### Electroacupuncture alleviates pain sensitivity and depression-like behavioural phenotypes in a bone cancer pain model

Bone cancer pain was modelled by injecting Lewis lung cancer cells into the left femoral cavity. Electroacupuncture treatment was started on the 10th day of cell injection lasting for 11 days. Electroacupuncture points were ST36, PC6 on the affected side.The parameters of the electro-acupuncture are 2HZ, 1mA, 20min (Fig 1A). The femur was dissected on day 21 after modelling to observe femoral changes. The model group showed bone destruction in the distal femur (Fig 1B), confirming successful modelling. Mice with similar baseline mechanical thresholds were selected for further modelling. The results showed a significant decrease in the threshold (Fig 1C). EA of ST36 acupoints significantly improved dysfunction, differing from the model group on day 14, and continued to improve in the EA group until day 21. In addition, EA increased sugar-water consumption in the model mice (Fig 1D). The above results support the analgesic effect of EA on inflammatory pain and the ameliorative effect on depression.

**Fig 1.**
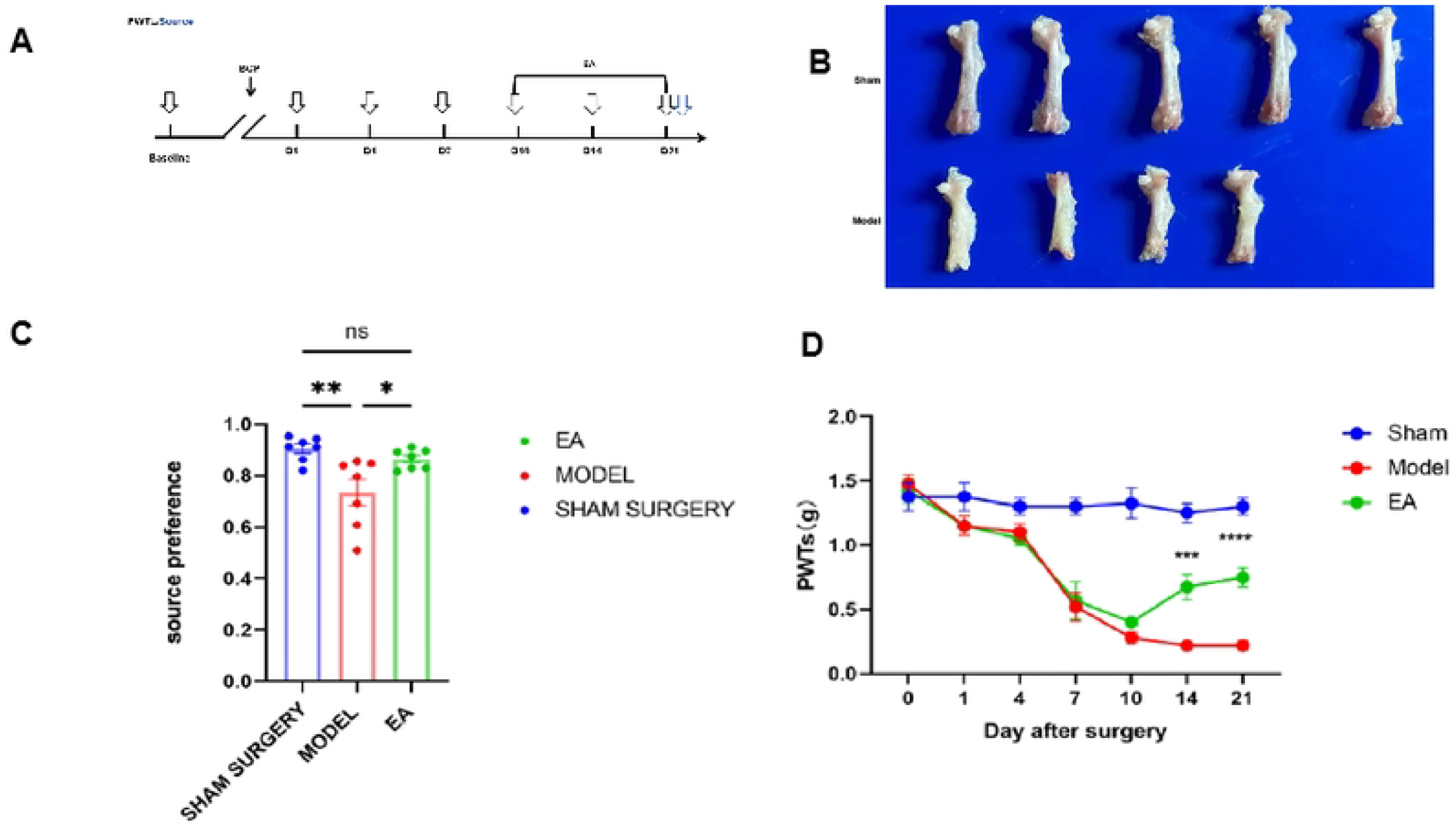
Source preference and Paw Withdrawal Mechanical Threshold (PWMT). EA improves pain threshold and depression in BCP mice. (A) Schedule of EA treatment and behavioural testing. Male C57 mice were injected with Lewis lung cancer cells in the femoral cavity on day 0 and then treated with EA (2 Hz, 1 mA, 20 min). and mechanically thresholded with von Frey. (B) Apparent valuation of distal femur destruction in the model group of mice. (C) Mechanical thresholds were detected using von Frey fibrin.EA significantly reversed the trend of decreased mechanical thresholds in model mice. There was no statistically significant difference at baseline between the three groups: sham group: 1.38*±*0.31 Model group: 1.48*±*0.21; Model+EA group: 1.43*±*0.25 One-way ANOVA with Bonferroni’s test F(12, 147)=10.07, P*<*0.0001. N=8 mice per group. * For model and model+EA groups. (D) Decreased glycogen consumption in the model group and increased glycogen consumption in the EA group, N=8 mice per group. (**P*<*0.01 vs. model group, **P*<*0.01,****P*<*0.0001.).

### Electroacupuncture modulates the STING/NF-*κ*B signalling pathway

STING is thought to be inextricably linked to the mechanisms of inflammation and involved in the modulation of pain. In this context, we tested whether the STING signalling pathway in BLA is associated with EA-mediated analgesia. We observed by (WB) a decrease in STING pathway protein levels in mice from the EA group (Fig 2A). a decrease in P-TBK1, TBK1, P-NF-*κ*B, and NF-*κ*B protein levels (Figs 2B-F).

**Fig 2.**
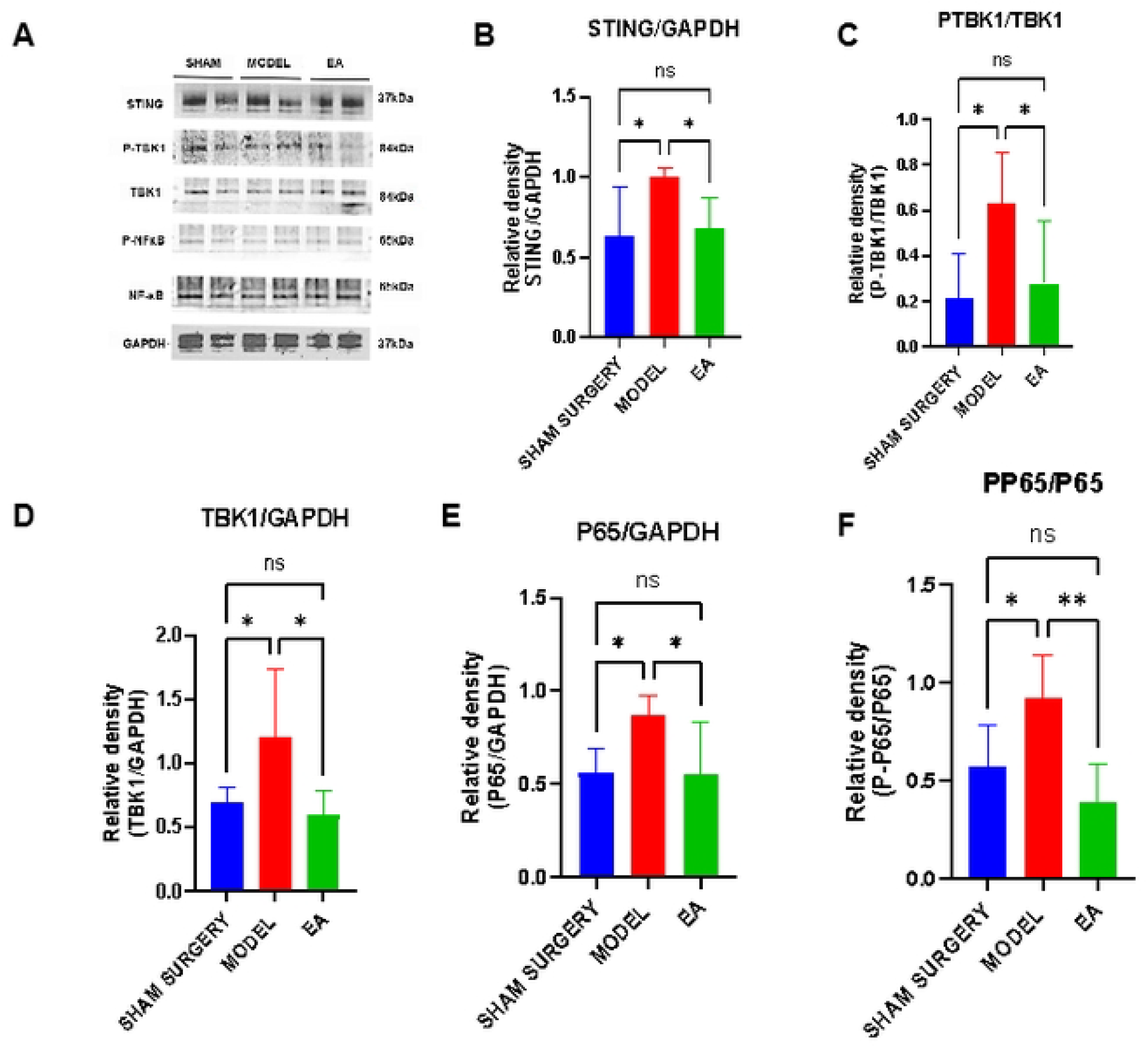
Western blot analysis of STING/NF-*κ*B in BLA. Western blot analysis of STING, TBK-1, P-TBK1, NF-*κ*B, p-NF-*κ*B in the BLA. EA depress the increase of STING, TBK-1, P-TBK1, NF-*κ*B, p-NF-*κ*B (A). Densitometric analysis results are presented as a ratio of protein intensities to GAPDH (B). Data (n = 6) are presented as means *±* SEM. *p *<* 0.01, **p *<* 0.05 vs model.

### Electroacupuncture inhibits microglia activation and inflammatory factor release

Immunofluorescence staining to study the activation of microglia in bone cancer pain. (Fig 3A). Immunofluorescence staining showed that the branch length and branch points of Iba-1 were increased in both sham and model+EA groups (Figs 3B-C). To directly explore the relationship with BCP microglia, we detected the pro-inflammatory factor IL-1*β* by enzyme-linked immunosorbent assay (ELISA), and the levels of this factor were significantly increased in pain mice, whereas the levels were significantly increased in the EA group (Fig 3D).

**Fig 3.**
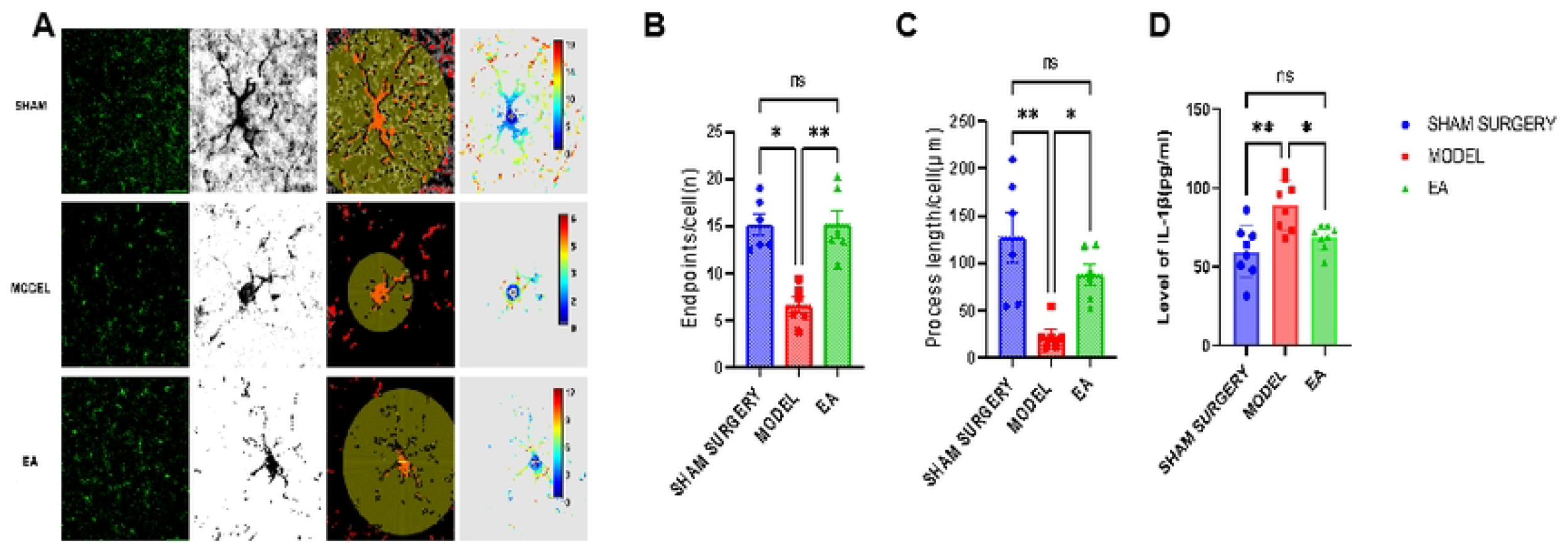
Immunofluorescence of IBA-1 and ELISA of IL-1*β* in BLA. Representative photos of immunofluorescence labeling of Iba-1 and ELISA of interleukin-1*β* (IL-1*β*) in the BLA (A). EA alletive the branch lengthes and branch points of Iba-1 and IL-1*β* expression (B-D). Scale bar = 100 *µ*m. Data (n = 6) are presented as means *±* S.E.M. *p *<* 0.01, **p *<* 0.001 vs model.).

## Discussion

In this study, we demonstrate how EA regulates bone cancer pain after LLC femoral inoculation. We found that EA modulates cancer pain and depression through the STING signalling pathway Consistent with a previous study [14], since STING modulates NF-*κ*B followed by the production of inflammatory factors, an increase in inflammatory factors leads to increased pain sensitivity. Significant relief of bone cancer pain on day 14 of treatment with EA. Notably, the treatment effect persisted on day 21, and microglia activation was also significantly inhibited, along with a decrease in inflammatory factors.

It was previously thought that neuroinflammation was caused exclusively by immune cells in the brain [15]. A previous study showed a time-dependent increase in STING expression in a BCP model [16]. Inhibition of STING resulted in significant pain relief.STING is a highly controversial protein. Some studies have used DMXAA (STING agonist) for pain relief in bone cancer pain models (days 3-7 after tumour inoculation), whereas our study focuses on the function of STING in mid-to late-stage tumours (day 10 after tumour inoculation).The different functions of STING may be related to the tumour stage, the different animal models.

Our study revealed that STING and its downstream signalling pathway proteins such as P-TBK1, TBK1 and P-NF-*κ*B, NF-*κ*B were up-regulated in the BLA brain region of BCP mice, along with an increase in pro-inflammatory cytokines (IL-1*β*), which may contribute to the development of advanced BCP in bone metastases. In addition, EA significantly reduced the upregulation of P-TBK1, TBK1 and P-NF-*κ*B, NF-*κ*B in the BLA of BCP mice. A close relationship between STING activation-induced hyperalgesia and inflammatory response was also found [17].

Activation of microglia in the BCP model has been well documented and the results showed a significant increase in Iba-1 expression in the BLA [18]. Zhang et al. reported that STING in mPFC may be a key regulator promoting microglia polarisation to the M1 phenotype [19]. Morphological changes after microglia injury have been reported in previous studies [20], and the present study refers to this literature for statistical methods [21]. Resting state microglia are highly branched with tertiary or quaternary branching structures. When inflammation, infection or other neurological diseases occur, microglia are rapidly activated. Activated microglia have enlarged cytosol, shortened protrusions, and rounded or rod-shaped cell morphology; activated microglia are further activated and restructured, with disappearing protrusions, amoeboid cell morphology and phagocytosis [22]. Morphological changes in microglia reflect the activation status of microglia, which is closely related to the severity of damage in the brain. There is a large body of literature demonstrating that after injury, microglials, in addition to changes in morphology, secrete a number of inflammatory factors into IL-1*β*, TNF-*α*, IL6, and so on, and these inflammatory factors seem to be able to have some effect on microglial morphology as well [23] [24]. For example, LPS is a common model used to induce inflammation in vivo, and the use of LPS can cause changes in microglial morphology and induce them to secrete inflammatory factors. After activation, microglials differentiate into two types, pro-inflammatory M1 and anti-inflammatory M2, and M1 microglials secrete a large number of inflammatory factors, such as TNF-*α*, IL-*β*, INF-*γ*, NO, and surface-expression of CD86, 68, etc. M2 microglials secrete IL-4, arignase1, Ym1, CD206, and IL-1, etc [25]. In response to injury, not only microglials react, but also the surrounding monocytes, which can be a major source of inflammation. When the brain is damaged, not only microglial cells respond, but also surrounding monocytes sometimes infiltrate into the brain, a process that depends on important factors such as CCR2, Ly-6C, and CX3CR1 [26]. They enter the brain and change their morphology, and interestingly, they look very much like microglial cells after the transformation [27].

Although our in vivo data confirm the critical role of STING in bone cancer pain, there are some limitations to our study. For example, although gender plays a crucial role in brain disease, we used only male rats. Therefore, there is a need to further explore the impact of these sex differences on the prognosis of bone cancer pain with depression; we will investigate whether these differences lead to changes in the STING pathway and microglia polarisation during bone cancer pain.

## Conclusion

Our results suggest that activation of STING in the BLA and proinflammatory factors downstream of STING lead to neuroinflammation and peripheral pain sensitisation in the BCP model.Activation of the STING pathway may be associated with microglia polarisation. Therapeutic strategies targeting this pathway may be a way to prevent or treat BCP.

## Acknowledgments

We would like to thank Professor Yihan He and Nenggui Xu for their contribution to this work, and we are particularly grateful to them for their edits and revisions. We declare that all authors have given their consent to the submission of this manuscript.

